# How do biases in sex ratio and disease characteristics affect the spread of sexually transmitted infections?

**DOI:** 10.1101/2020.06.30.179747

**Authors:** Naerhulan Halimubieke, Alistair Pirrie, Tamás Székely, Ben Ashby

## Abstract

The epidemiology of sexually transmitted infections (STIs) is inherently linked to host mating dynamics. Studies across many taxa show that adult sex ratio, a major determinant of host mating dynamics, is often skewed - sometimes strongly - toward males or females. However, few predictions exist for the effects of skewed sex ratio on STI epidemiology, and none when coupled with sex biased disease characteristics. Here we use mathematical modelling to examine how interactions between sex ratio and disease characteristics affect STI prevalence in males and females. Notably, we find that while overall disease prevalence peaks at equal sex ratios, prevalence per sex peaks at skewed sex ratios. Furthermore, disease characteristics, sex-biased or not, drive predictable differences in male and female STI prevalence as sex ratio varies, with higher transmission and lower virulence generally increasing differences between the sexes for a given sex ratio. These findings may be due to a balance between increased per-capita mating in the less common sex, and a reduction in mating rate - hence disease prevalence - at the population level. Our work reveals new insights into how STI prevalence in males and females depends on a complex interaction between host population sex ratio and disease characteristics.

## 1. Introduction

Sexually transmitted infections (STIs) – defined as any pathogen that is transmitted during copulation – are ubiquitous in the animal kingdom, often causing chronic infections with low recovery rates and reduced reproductive success, even sterility [1, 2]. STIs typically exhibit contrasting epidemiological dynamics to non-STIs, with transmission likely to be frequency-rather than density-dependent (i.e. the number of sexual contacts per capita is invariant to population size). Another major difference is that STIs are primarily transmitted between the sexes (especially in non-human populations), which creates a natural bipartite network where the population is split into two disjoint sets (males and females) and connections only exist between individuals from distinct sets. In contrast, non-STI transmission networks are generally not bipartite. Sex ratio, which can vary greatly in non-human populations [3-6], will therefore likely have a profound impact on STI epidemiology compared to non-STI epidemiology, as it disrupts the structure of the contact network [7, 8]. There is growing evidence shedding light on the relationship between host mating system structure and STI dynamics [9-11], much of which focuses on human populations [12, 13]. However, the implications of variation in sex ratio for STI epidemiology are not well understood.

Evidence has shown that sex ratio has a crucial role in mating system variation in wild populations [5, 14]. Sex ratio can vary at different stages of the life cycle, with the ratios of males to females at conception, at birth and during adult life (termed primary, secondary, and tertiary or adult sex ratio, respectively). Recent studies show that adult sex ratio may deviate from 1:1 in a variety of organisms: butterflies, reptiles and birds often have male-biased sex ratios with up to 90% of the population being male, whereas female-biased adult sex ratios are common in insects and mammals, with up to 100 females for every male [3-6]. Variation in the adult sex ratio can alter the mating opportunities of breeding males and females and may select for different mating systems [15, 16], the latter has significant consequences for STI dynamics [17-20]. Interestingly, *Wolbachia* bacteria (transmitted vertically rather than sexually in insect hosts) are known to distort the birth sex ratio, as it is maternally inherited and kills male embryos, consequently, disrupt the mating dynamics of their hosts [7, 21, 22].

Adult sex ratio is clearly crucial for mating dynamics and hence STI transmission, but it is shaped by multiple factors, including primary and secondary sex ratio as well as differential survival during juvenile and adult stages [23]. STIs may cause differential mortality between the sexes, potentially increasing or decreasing the ratio of adult males to females, causing feedback between STI transmission and sex ratio [5]. In general, theory predicts that the rarer sex in the population should exhibit higher STI prevalence since they have a higher per capita mating rate, all else being equal [8]. For example, two-spot ladybirds *Adalia bipunctata* exhibit male-biased patterns of STI prevalence in female-skewed populations [8]. A follow-up experimental study by Pastok et al. [24] established that the presence of male-killing bacteria at high prevalence skews the sex ratio towards a female-biased population, and results in male-biased STI prevalence. Together, these studies suggest that there are likely to be complex interactions between STIs and adult sex ratios. However, the extent to which STI prevalence in males and females depends on sex ratio and disease characteristics, which may vary between the sexes, has yet to be explored.

Many pathogens – both STIs and non-STIs - show sex bias in disease characteristics (e.g. transmissibility, virulence), which can influence disease dynamics [25, 26]. Two main hypotheses have been proposed to explain sex-biased disease characteristics. The *physiological hypothesis* emphasises that the interactions between sex hormones and the immune system render one sex more susceptible to infection and disease [27]. For example, in mammals, males have consistently weaker immune competence than females, and this correlates with male-biased disease prevalence, mortality, and female-biased adult sex ratio [28-30] On the other hand, the *behavioural hypothesis* posits that sexual differences in behaviour may cause sex-specific exposure to pathogens [31]. For instance, infections by arthropods, helminths and unicellular parasites are often male-biased in mammals but not in birds [32]. Sex differences are expected, since males tend to be more active in reproductive behaviours such as combat for females, territorial defence and foraging, due to more intense sexual selection in males than females, therefore increasing the chance of exposure to pathogens. In human populations, gender-related behavioural differences may render one sex more exposed to certain contagions (STIs or non-STIs). For example, Ebola haemorrhagic fever outbreak that occurred in the Congo and Gabon in 2001-2002 suggested more men than women were infected during the early stages of the outbreak because men spent more time working away from home, where they were more likely to come into contact with wild animals [33]. Regardless of whether sex biases in disease characteristics are due to physiological or behavioural differences between males and females, there are likely to be important implications for variation in disease prevalence between the sexes.

In the context of STIs, current empirical research in sex-biased disease characteristics is mainly focused on humans. Some studies have established that there is a higher risk of transmission of certain STIs from men to women than vice versa due to STIs being present in ejaculate [34, 35]; another study shows that human T-lymphotropic virus exhibits male-biased virulence, potentially because transmissions occur during pregnancy, birth or breast-feeding, and so selection has favoured lower virulence in women [36]. Empirical studies of sex-biased STI characteristics in animals are limited, but mathematical models have shown how sex-biased disease characteristics influence transmission dynamics (reviewed in [11], also see [12]). However, the interplay between sex ratio and disease-characteristics (especially when these are sex-biased) and their effects on STI prevalence have yet to be investigated. Here we use mathematical modelling to examine the effects of biases in population sex ratio and disease characteristics on STI epidemiology. Specifically, we investigate how (1) variation in population sex ratio; (2) sex-biases in transmission and disease-associated mortality (virulence) in adults; and (3) the interactions between sex ratio and disease characteristics, impact on STI prevalence in males and females.

## 2. Methods

We model the dynamics of an STI in a randomly mixing population. The epidemiological and mating dynamics are described by the following system of ordinary differential equations:

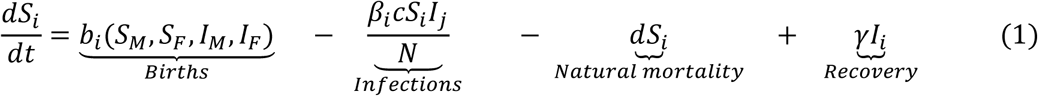

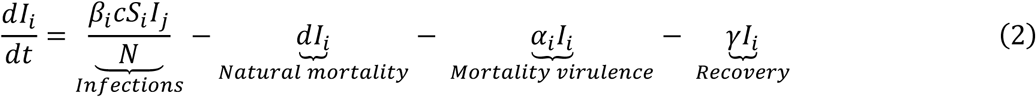

where *i, j* ∈ {*M, F*}, *i* ≠*j*, correspond to males and females. The birth rates for each sex are

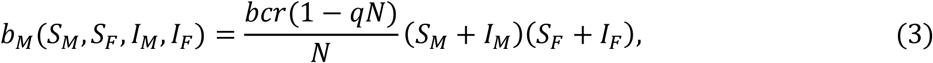

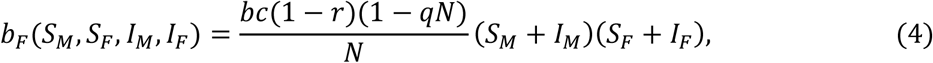

and: *S*_*i*_ and *I*_*i*_ (equivalently, *S*_*j*_ or *I*_*j*_), are the number of susceptible and infected individuals, respectively, in each sex; *N* = *S*_*M*_ + *S*_*F*_ + *I*_*M*_ + *I*_*F*_ is the total population size; *r* is the sex ratio at birth (the proportion of offspring that are male), with 0 ≤ *r* ≤ 1; *c* ≥ 0 is the pairwise rate at which males and females mate given frequency-dependent contact; *q* represents the strength of density-dependence on births; *d* is the natural mortality rate; *α*_*i*_ is the disease-associated mortality rate (virulence) in sex *i*; *β*_*i*_ is the transmission rate from sex *j* to sex *i* per sexual contact; *γ* is the recovery rate. The probability that a susceptible female mates with an infected male (similarly, for the reverse scenario) is equal to 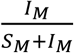, which is obtained by dividing the mating rate with infected males 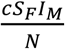 by the total mating rate for susceptible females 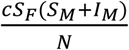.

For simplicity, we assume that: (i) both sexes have the same recovery rate from the STI; (ii) both sexes have the same natural mortality rate; (iii) the per-capita mating rate is frequency-dependent, which means that larger populations do not have a higher per-capita mating rate than smaller populations; (iv) there is no structuring or choice in the mating system (mating is random); (v) there is no juvenile period; and (vi) there is no effect of parental care on the population dynamics.

The disease-free equilibrium 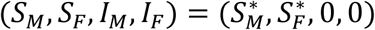, of this system occurs at:

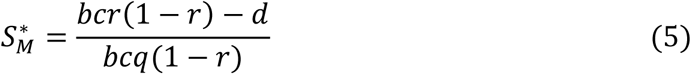

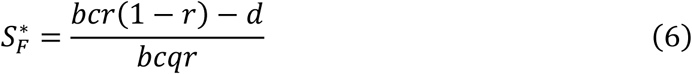

A newly introduced STI will spread in a susceptible population when the basic reproductive ratio, *R*_0_ is greater than 1 (*R*_0_ is calculated as the maximum eigenvalue of the Jacobian of the system, evaluated at the disease-free equilibrium), where:

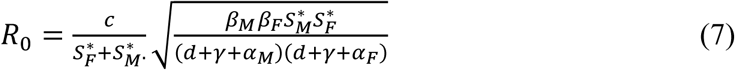

In order to guarantee that *R*_0_ > 1, we require that 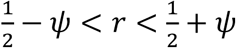, where 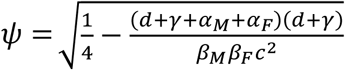. To test the interactions between sex ratio and STI prevalence in males and females, we start with no sex bias in STI characteristics. This means that males and females are equally susceptible to the pathogen and the mortality caused by the pathogen is same in both sexes (*α*_*M*_ =*α*_*F*_=*α; β*_*M*_ =*β*_*F*_=*β*). STI prevalence in each sex is given as the number of infected individuals in that sex divided by the total number of individuals in that sex (at equilibrium, as indicated by asterisks):

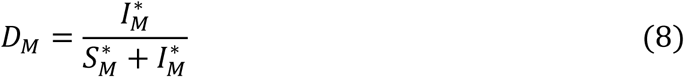

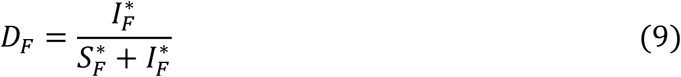

To examine the impact of sex-biased disease characteristics on the prevalence of STIs in males and females, we fix female susceptibility (*β*_*F*_) and disease-related mortality (*α*_*F*_) and vary the corresponding parameters in males. The model is symmetric up to labelling, so the results are analogous for the converse scenario. We primarily use numerical analysis to determine disease prevalence as the endemic equilibrium of equations 1-2 is mostly intractable to algebraic analysis.

## 3. Results

### 3.1 Non-sex-biased disease characteristics

We begin by exploring how sex ratio impacts on STI prevalence in males and females when there are no sex-biases in disease characteristics (i.e. *α*_*M*_ =*α*_*F*_=*α; β*_*M*_ =*β*_*F*_=*β*). Consistent with previous theoretical and empirical work [7, 8], the less common sex exhibits higher STI prevalence at equilibrium (Fig. 1). Intuitively, when there are no sex biases in disease characteristics, the overall prevalence of the STI is always maximised at an equal sex ratio (*r* = 0.5). However, both the extent to which STI prevalence differs between the sexes (i.e. the difference between the male and female curves in Fig. 1a-b, shown in Fig. 1c-d) and the point at which STI prevalence peaks in each sex (i.e. the sex ratio that maximises the male and female curves, indicated by circles in Fig.1a-b) depends on the transmission rate and virulence. Specifically, as the transmission rate increases or as the virulence decreases, the difference between male and female STI prevalence for a given sex ratio tends to increase, especially for more extreme sex ratios (Fig. 1c-d). Furthermore, for higher transmission rates/lower virulence, the sex ratio at birth for which disease prevalence peaks becomes more extreme. Thus, while overall STI prevalence is qualitatively unchanged by variation in the transmission rate and virulence (i.e. it always peaks at *r* = 0.5), STI prevalence per sex is highly sensitive to disease characteristics.

**Figure 1.**
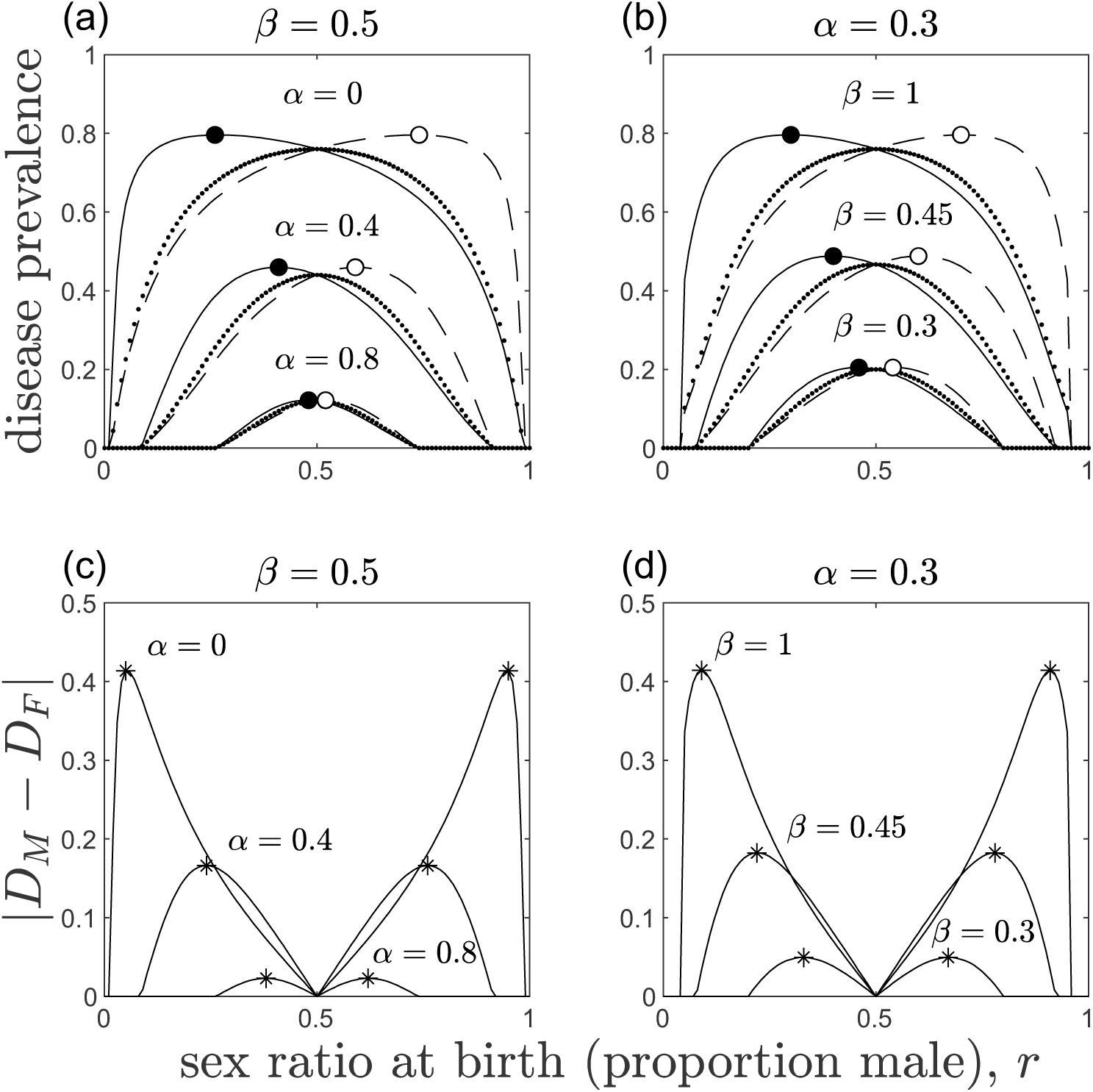
Effects of sex ratio at birth (proportion male), *r*, on STI prevalence when there are no sex biases in disease characteristics: (a, c) variation in disease-related mortality, *α*_*M*_ = *α*_*F*_ = *α*; (b, d) variation in transmission probability, *β*_*M*_ = *β*_*F*_ = *β*. As disease-related mortality increases or the transmission probability decreases, overall STI prevalence decreases, and the difference between male and female STI prevalence for a given sex ratio decreases. (a, b) STI prevalence in males (solid line), females (dashed line), and overall (dotted line). Markers show where STI prevalence peaks in males (black) and females (white). (c, d) Absolute difference in STI prevalence between males (*D*_*M*_) and females (*D*_*F*_), corresponding to the absolute difference between the solid and dashed lines in (a) & (b), with markers showing where the absolute difference peaks. Other parameters are held constant throughout to ensure *R*_*0*_ is greater than 1: *δ* = 0.1; *γ* = 0.2; *c* = 5; *b* = 1; *q* = 0.001.

We can see this analytically for variation in the transmission rate if we make the following simplifying assumptions: (1) there is no disease-associated mortality (*α*_*M*_ = *α*_*F*_ = 0); (2) the disease transmission rates are equal (*β*_*M*_ = *β*_*F*_ = *β*); and (3) there is no recovery (*γ* = 0). These assumptions ensure that the adult sex ratio is equal to the sex ratio at birth, which greatly simplifies the analysis. First, we must calculate the equilibrium population size, *N*\*. Let *M* = *S*_*M*_ + *I*_*M*_ be the number of males in the population and *F* = *S*_*F*_ + *I*_*F*_, be the number of females in the population. Then at equilibrium, *M* = *sN*\*, and *F* = (1 − *r*)*N*\*. Now, 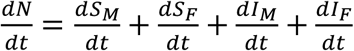, and evaluating this gives

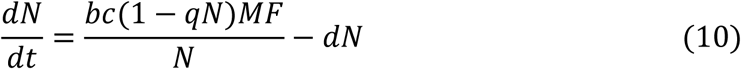

Substituting in *M* = *sN*\* and *F* = (1 − *r*)*N*\*, and solving 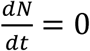, the resulting (non-trivial) equilibrium population size is:

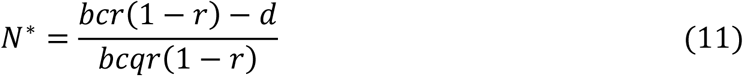

We can now substitute the equilibrium population size into the full system of equations, giving the endemic equilibrium:

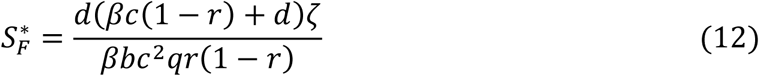

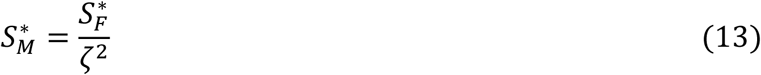

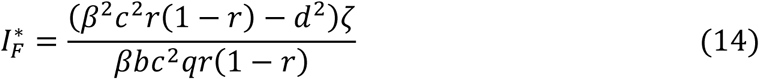

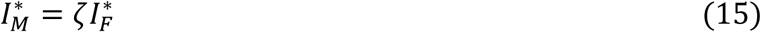

where 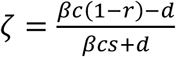. The disease prevalence in males and females is therefore

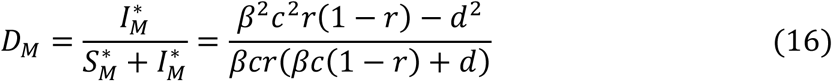

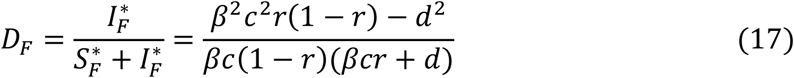

To find the peak in disease prevalence with respect to *r*, one must differentiate equations 16-17 with respect to *r* and then find where they are equal to zero, i.e., find 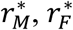 such that 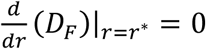 and 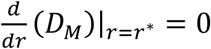. For females, STI prevalence peaks at

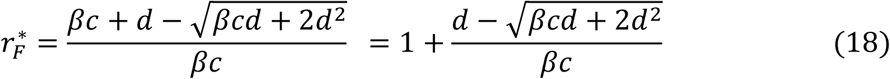

and for males STI prevalence peaks at

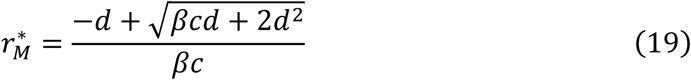

(Note that the other root lies outside *r* ∈ [0,1]). Taking derivatives with respect to *β*, we find that 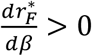 and 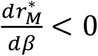. Thus, as the transmission rate, *β*, increases, the sex ratio at which STI prevalence peaks in each sex always becomes more extreme.

### 3.2 Sex-biased disease characteristics

We now consider the effects of sex-biased disease characteristics on STI prevalence. Specifically, we investigate how STI prevalence (per sex and overall) varies when there are biases in (1) female-to-male (*β*_*M*_) or male-to-female (*β*_*F*_) transmission, and/or (2) disease-associated mortality (virulence) for males (*α*_*M*_) or females (*α*_*F*_). Since the only differences in the model between males and females are these parameters, without loss of generality we fix *β*_*F*_ and *α*_*F*_ whereas vary *β*_*M*_ and *α*_*M*_ accordingly.

Sex differences in transmission rate and in virulence influence disease prevalence in both sexes, although the extent of their influence on males and females depends on sex ratio. A higher female-to-male transmission rate relative to the male-to-female transmission rate raises disease prevalence in both sexes, although interestingly the influence of sex ratio differs between the sexes. With greater female-to-male transmission, the disease prevalence peak shifts toward male-biased sex ratios in females whereas for males there is relatively little change (e.g. Fig. 2a-c; Fig. 3a-c). Similarly, lower disease-associated mortality in males relative to females also universally increases disease prevalence and shifts the peaks of the disease prevalence curves. However, lower values of *α*_*M*_ generally cause the disease prevalence peaks in each sex to shift towards lower values of *r*, with greater effects on male than female STI prevalence (e.g. Fig. 2b, e, h; Fig. 3d-f).

**Figure 2.**
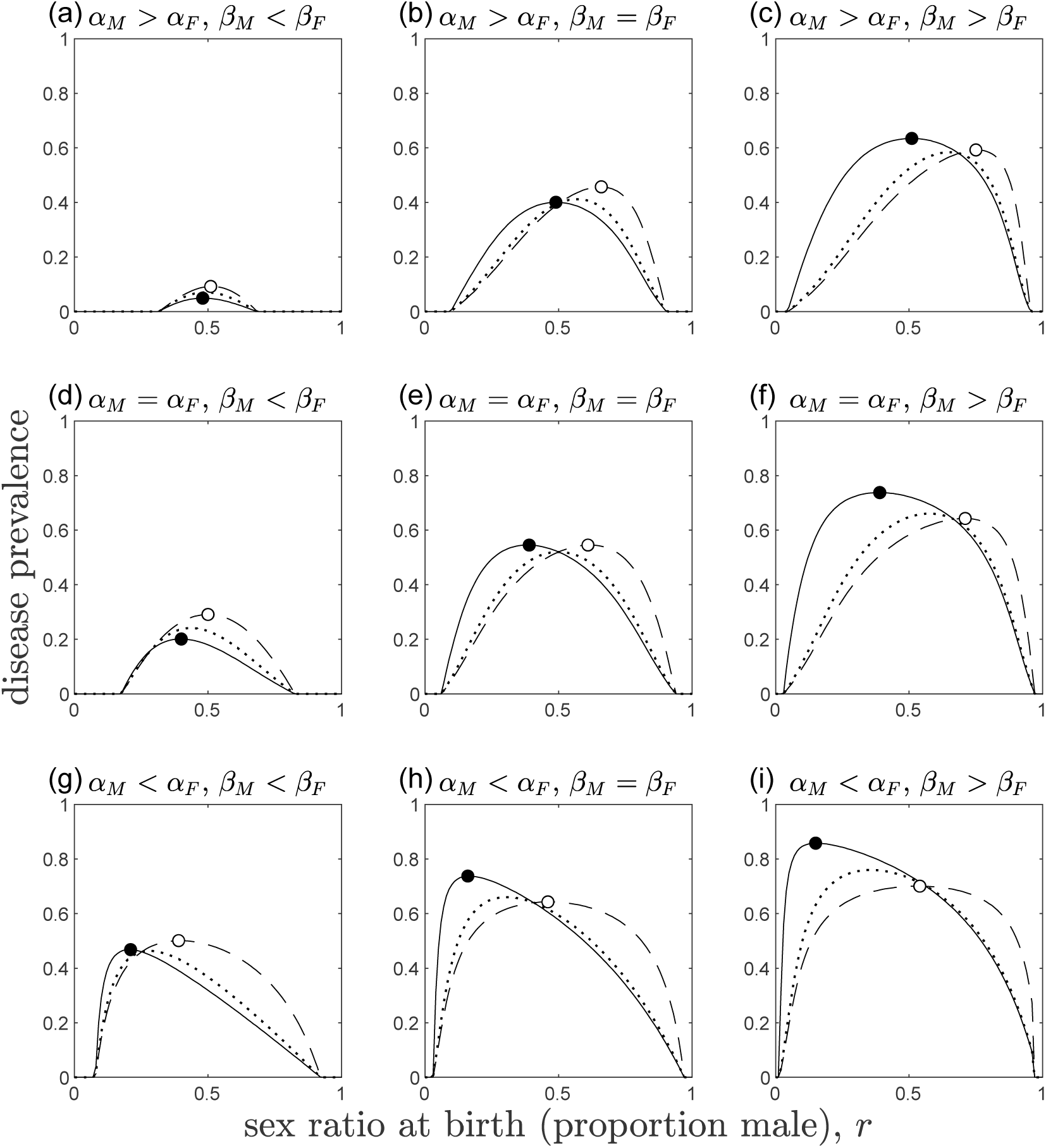
STI prevalence in males (solid line), females (dashed line), and overall (dotted line) as a function of the sex ratio at birth (proportion male), *r*, when there are sex biases in STI transmission rate and virulence. The parameters *β*_*F*_ = 0.5 and *α*_*F*_ = 0.3 are held constant throughout, while *β*_*M*_ ∈ {0.2,0.5,1} and *α*_*M*_ ∈ {0,0.3,0.6} vary between panels. Other parameters as described in Fig. 1.

**Figure 3.**
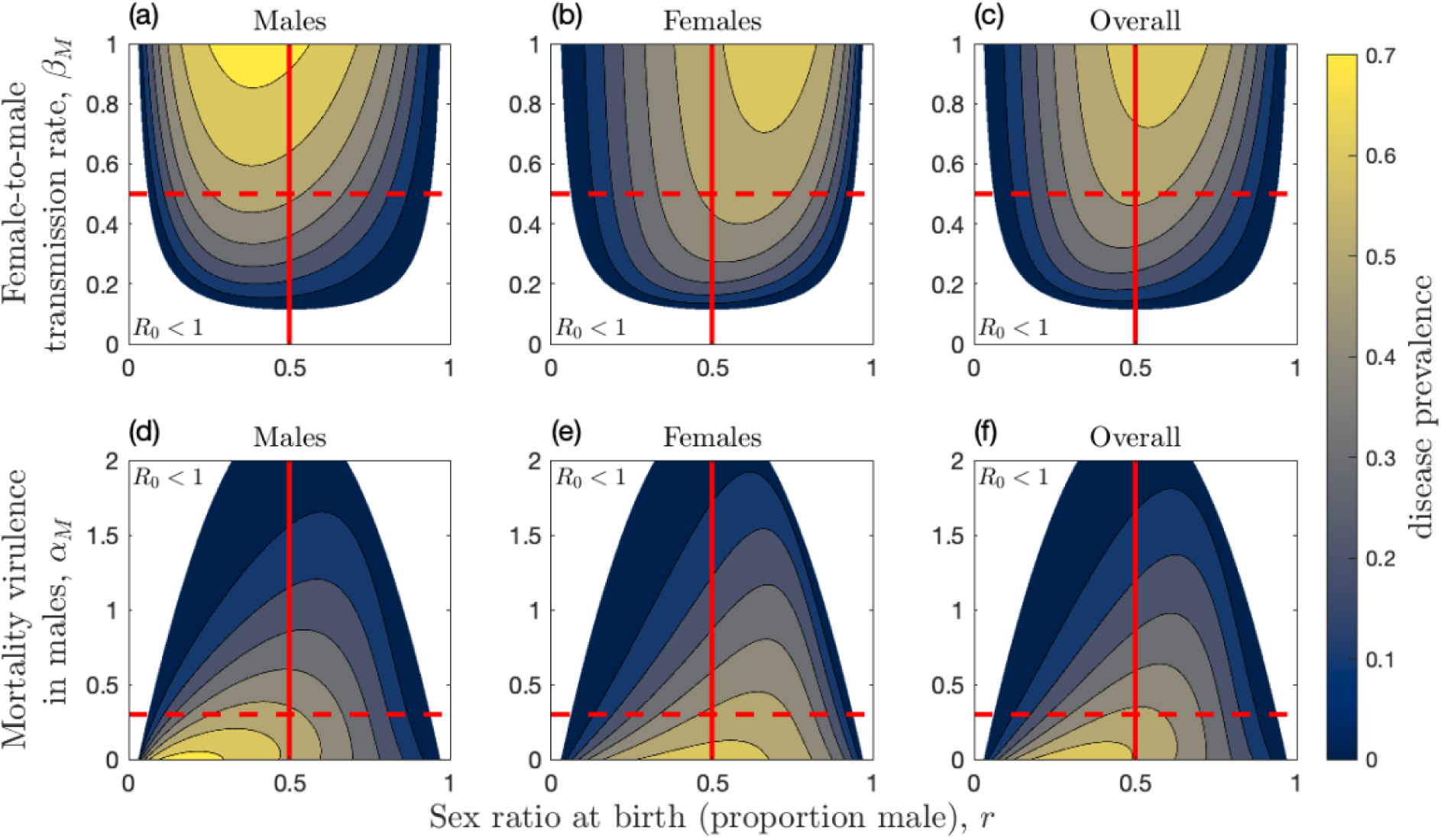
STI prevalence in (a, d) males, (b, e) females, and (c, f) overall as a function of the sex ratio at birth and: (a-c) female-to-male transmission rate (*β*_*M*_); (d-f) mortality virulence in males (*α*_*M*_). The red lines show when the sex ratio at birth is equal (solid) and when there is no sex bias in disease characteristics (dashed): (a-c) *β*_*M*_ = *β*_*F*_ = 0.5; (d-f) *α*_*M*_ = *α*_*F*_ = 0.3. The white region in each panel corresponds to conditions where the STI is unable to persist (*R*_0_ < 1). The parameters *β*_*F*_ = 0.5 and *α*_*F*_ = 0.3 are held constant throughout, Other parameters as described in Fig. 1.

We see consistent patterns when transmission and virulence are varied simultaneously (Fig. 4). By measuring the sex ratio at birth, *r*, for which disease prevalence peaks, we analyse how the interaction between transmission, virulence, and sex ratio leads to changes in disease prevalence. When disease-associated mortality in males is relatively low (*α*_*M*_ < *α*_*F*_), variation in the female-to-male transmission rate (*β*_*M*_) has little impact on the sex ratio where disease prevalence peaks in males (Fig. 4a) and overall (Fig. 4c), but the effect in females is much more pronounced, with the skew generally increasing for greater *β*_*M*_ (Fig. 4b). When disease-associated mortality in males is relatively high (*α*_*M*_ > *α*_*F*_), the sex ratio at birth where disease prevalence peaks in males and overall also increases with *β*_*M*_, but to a lesser extent than in females. It is also clear from the contours in Fig. 4 that variation in either transmission or virulence can cause non-monotonic changes in the disease prevalence curves. For example, the horizontal and vertical red lines in Fig. 4 (indicating *β*_*M*_ = *β*_*F*_ or *α*_*M*_ = *α*_*F*_) intersect some of the contours twice, which means that varying either *α*_*M*_ or *β*_*M*_ can cause a non-monotonic change in the skew of the disease prevalence curves. Together, these results show that interactions between sex-biased disease characteristics and sex ratio can lead to major changes in STI prevalence within each sex and overall, and that the differences between male and female STI prevalence can be exacerbated by greater transmission rates or lower mortality rates, in one or both sexes.

**Figure 4.**
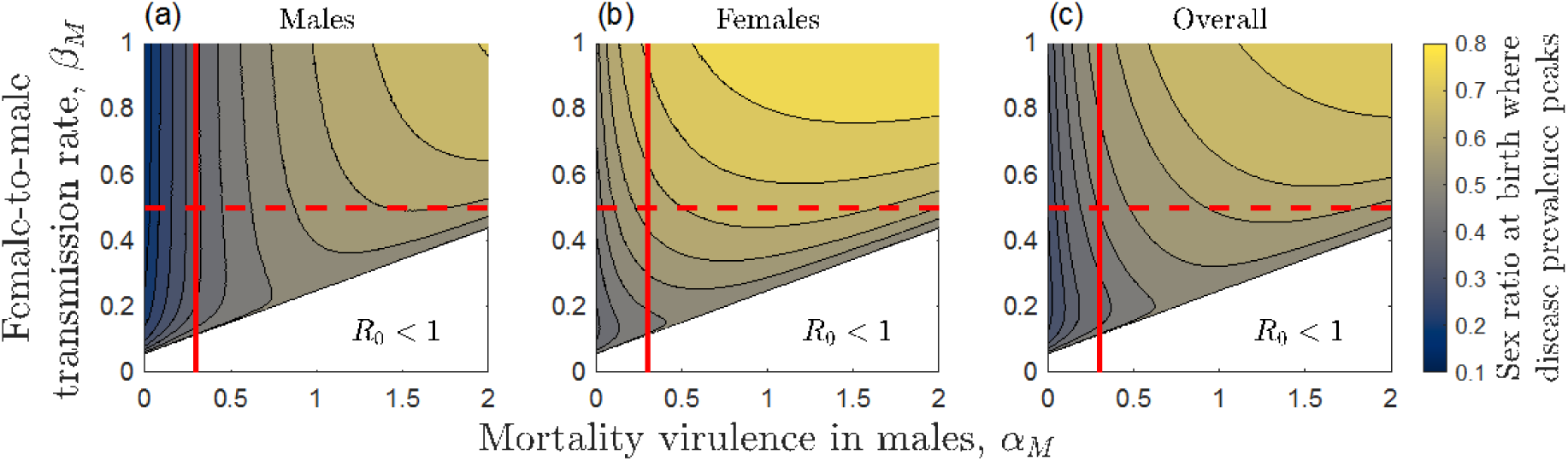
The sex ratio at birth for which disease prevalence peaks in: (a) males, (b) females, and (c) overall, as a function of disease-associated mortality for males (*α*_*M*_) and female-to-male transmission rate (*β*_*M*_). The red lines show when there is no sex bias in transmission (*β*_*M*_ =*β*_*F*_ =0.5; dashed) and when there is no sex bias in virulence (*α*_*M*_=*α*_*F*_=0.3; solid). The white region in the bottom right corner of each panel corresponds to conditions where the STI is unable to persist (*R*_0_ < 1). Other parameters as described in Fig. 1.

## 4. Discussion

Studies of STI epidemiology emphasise the importance of the host mating system for STI dynamics [8-11, 19, 24]. Since sex ratio influences mating systems, here we used mathematical modelling to examine the interaction between sex ratio and disease-characteristics, and their effects on STI prevalence in males and females. At a fundamental level, our results reveal how sex differences in STI prevalence within and between populations may be explained by both variation in sex ratio and disease characteristics. Moreover, our model shows that disease-characteristics, whether they be sex-biased or not, can increase or decrease differences in male and female STI prevalence.

In a population with a skewed adult sex ratio, the less common sex has more prospective mates and a higher per-capita mating rate than the more common sex. This pattern can be seen in human and non-human societies; for instance, in human communities with a male-skewed adult sex ratio, men are more likely to purchase sex and women are more likely to have multiple sex partners [37, 38]. Consistently, in bird populations mating opportunities are related to adult sex ratios: in male-biased adult sex ratio, females have higher mating opportunities than males, whereas in female-skewed populations males re-mate faster than the females [39-41]. Hence, with high variance in mating success, many individuals of the more common sex may remain unmated or have a low mating rate, whereas the less common sex has a higher mating rate and hence greater exposure to infectious partners. Thus, equilibrium disease prevalence in the more common sex is expected to be lower than in the rarer sex, consistent with previous theoretical and empirical studies [7, 8, 10, 11].

Our model shows how STI prevalence – overall and in each sex – depends on both sex ratio and disease characteristics. We have shown that even when there are no sex biases in disease characteristics, increasing the transmission rate or decreasing mortality virulence generally leads to a greater difference in STI prevalence between the sexes for a given sex ratio (Fig. 1c-d). We have also shown that the sex ratio for which disease prevalence peaks in each sex depends on disease characteristics, again even when the disease itself shows no sex biases in transmission or virulence. The reason for these patterns can be understood in terms of a balance between an increase in the per-capita mating rate for the less common sex and a decrease in overall disease prevalence as the sex ratio becomes more skewed. Suppose the adult sex ratio is initially equal and that there are no biases in disease characteristics, hence equilibrium STI prevalence is the same in both sexes. If we gradually remove males from the population (similarly for females), then the per-capita mating rate for the remaining males increases, and hence so does their risk of infection, at least initially. However, skewing the sex ratio in either direction reduces the overall mating rate, and therefore always lowers disease prevalence at the population level. As one continues to remove males from the population, overall disease prevalence begins to fall more sharply because uninfected males are increasingly less likely to mate with infected females. Thus, disease prevalence eventually begins to fall in males as well. Once the population crosses a critical sex ratio, *R*_0_ falls below 1 and the disease is driven extinct. Higher transmission and lower virulence both increase overall disease prevalence and therefore buffer against the effects of skewed sex ratios, which shifts the sex ratio at which disease prevalence peaks in each sex to more extreme values. This can be seen in Fig. 1a-b, where the gradient of the overall disease prevalence curves is flatter for higher values of *β* and lower values of *α*.

We also explored the effects of sex-biased transmission and virulence. Sex-biased disease characteristics are common in wild populations as well as in human populations [29, 42-44], and previous studies have looked into the behavioural and physiological causes of such disparities and their implications for population dynamics [26, 27, 31]. However, the interaction between these biases and STI prevalence within each sex have previously been overlooked. Our model suggests that sex-biased disease characteristics may independently influence STI prevalence in a similar way as the unbiased transmission rate or virulence (see above). However, the prevalence curves for each sex are no longer symmetrical when there are biased STI characteristics (Fig. 2). Our model also reveals that interactions between sex ratio and sex-biased disease characteristics can lead to differential and non-intuitive patterns for STI prevalence. For example, while increasing the bias in transmission from females to males causes STI prevalence in females to peak at a more extreme sex ratio, there are only marginal effects on males, yet the effects of male-biased virulence on disease prevalence are much more pronounced in males than females (as shown in Fig. 3). The model also predicts that increasing or decreasing sex-biased disease characteristics may lead to non-monotonic variation in STI prevalence. In the absence of mathematical modelling, it is unlikely that one would be able to intuit such effects, but our results suggest that there are likely to be complex interactions between sex-biased disease characteristics and population sex ratio that shapes STI epidemiological dynamics.

Our model is simple and makes few biological assumptions. Therefore, follow-up analyses are necessary to investigate biological factors mediating STI prevalence that are not captured in our model. First, previous studies have emphasised the importance of pair formation in STI prevalence [45, 46], however, our model assumes mating is random, there is no variance in mating rate, and hosts do not form pair-bonds [47-50]. This was important for the present study as we wanted to focus on effects of sex ratio and disease characteristics, all else being equal. However, future work would benefit from taking pair-formation into account, as a series of studies in human STIs have shown the variations in form and duration of partnerships may have an effect on pathogen prevalence [51-54]. Second, previous studies have shown how variance in mating rate [10, 11] and mate choice [49, 50, 55] affect disease dynamics, which in turn may influence host and STI evolution in non-intuitive ways. When one sex experiences higher variance in mating rate, the mating population is effectively sex-skewed and so we would expect this to generally have a similar effect on STI prevalence as seen in our model. If infection status impacts on mating rate [47, 49, 50, 56], then the effects may be more complex, especially if mate choice is driven by one sex.

Previous models have explored how STIs might affect the evolution of host mating dynamics [46-50], but the focus of the present study has been on STI prevalence. However, given that there are interactions between sex ratio, sex-biased disease characteristics and mating dynamics, there are likely to be evolutionary implications for these factors on hosts and STIs, which should be the target of future research. In particular, it is well-established that disease prevalence is important for the evolution of host defence [57], and so biases in sex ratio or disease characteristics are likely to influence selection for traits such as resistance, tolerance or mate choice through STI prevalence [49, 50]. Additionally, since females are more important than males for population growth in many species, female-biased disease prevalence might be more likely to increase population extinction risk.

Our results have several implications for STIs in natural populations. First, when sex ratios are skewed, we should expect STIs to be more prevalent in the rarer sex, although the relative prevalence of co-circulating STIs in males and females will differ due to variation in transmission and virulence (Fig. 1c-d). For example, in a female-skewed population, an STI with a high transmission rate is likely to have a higher prevalence in males relative to females compared to an STI with a low transmission rate. Second, our model predicts that variance in the sex ratio between populations to drive STI prevalence in males and females, with divergence from equal sex ratios generally increasing sex differences in prevalence (except at extreme sex ratios). Finally, we should expect the interaction between sex ratio and sex-biased transmission or virulence to consistently have a stronger effect on STI prevalence in one sex than in the other.

## 5. Conclusion

To conclude, the objective of our model was to examine the interplay between sex ratio and disease characteristics and explore their effects on STI prevalence. Our key message is that while the less common sex is predicted to exhibit higher STI prevalence at equilibrium, disease characteristics such as transmission rate and virulence – sex-biased or not – combined with sex ratio, drive differential patterns in male and female STI prevalence.

## Supporting information

Source code for the model and figures

## Author contributions

BA, NH and AP designed and analysed the model. NH and AP co-wrote the first draft, and all authors contributed to revisions of the manuscript.

## Data availability

Source code is available in the Supplemental Material and in the following Github repository: https://github.com/ecoevogroup/Halimubieke_et_al_2020.

