## Supplementary material for "How do biases in sex ratio and disease characteristics affect the spread of sexually transmitted infections?": Source code for the model and figures: fig1.m

```
% Fig 1


t_max = 1000;
alpha = 0.3;
beta = 0.5;

ALPHA = [0,0.4,0.8];
BETA = [0.3,0.45,1];

r = linspace(0,1,101);
plotflag = 0;
c = 5;
d = 0.1;
gamma = 0.2;
sub = 0;

labs = {'(a)','(b)','(c)','(d)'};

pos1 = [0.1,0.29,0.63];
pos2 = [0.63,0.29,0.12];

figure(1)
clf
set(gcf,'color','w')
for i1=1:2
    for j1=1:3
        if(i1==1)
            alphaM = ALPHA(j1);
            alphaF = ALPHA(j1);
            betaM = beta;
            betaF = beta;
        else
            alphaM = alpha;
            alphaF = alpha;
            betaM = BETA(j1);
            betaF = BETA(j1);
        end
        
        males=zeros(length(r),1);
        females=zeros(length(r),1);
        overall=zeros(length(r),1);
        diff1=zeros(length(r),1);
        for i=1:length(r)
            [t,x] = sexratio1(alphaM,alphaF,betaM,betaF,r(i),plotflag);
%             [t,x] = sexratio1_fast(t_max,alphaM,alphaF,betaM,betaF,d,d,gamma,gamma,b,c,1,1,q,r(i));
            if(sum(x(end,3:4),2)>1e-6)
                males(i,1) = x(end,3)/sum(x(end,[1,3]),2);
                females(i,1) = x(end,4)/sum(x(end,[2,4]),2);
                overall(i,1) = sum(x(end,3:4),2)/sum(x(end,:),2);
                diff1(i,1) = x(end,3)/sum(x(end,[1,3]),2) - x(end,4)/sum(x(end,[2,4]),2);
            end
        end
        maxMP = max(males);
        maxFP = max(females);
        peaks = [r(find(males==max(males),1)),r(find(females==max(females),1))];
        
        maxdiff1 = max(diff1);
        maxdiff2 = min(diff1);
        diffpeaks = [r(find(diff1==max(diff1),1)),r(find(diff1==min(diff1),1))];
        diff1 = abs(diff1);
        
        subplot(2,2,i1)
        hold on
        plot(r,males,'k')
        plot(r,females,'k--')
        plot(r,overall,'k.','markersize',4)
        plot(peaks(1),maxMP,'ko','MarkerSize', 5,'markerfacecolor','k')
        plot(peaks(2),maxFP,'ko','MarkerSize', 5,'markerfacecolor','w')
        ylim([0 1])
        if(i1==1)
            if(j1==1)
                text(0.42,maxMP+0.1,strcat('$\alpha=',num2str(alphaF),'$'),'interpreter','latex','fontsize',10)
            else
                text(0.4,maxMP+0.1,strcat('$\alpha=',num2str(alphaF),'$'),'interpreter','latex','fontsize',10)
            end
        else
            if(j1==3)
                text(0.42,maxMP+0.1,strcat('$\beta=',num2str(betaF),'$'),'interpreter','latex','fontsize',10)
            else
                text(0.4,maxMP+0.1,strcat('$\beta=',num2str(betaF),'$'),'interpreter','latex','fontsize',10)
            end
        end
        set(gca,'fontsize',8)
        box on
        set(gca,'ytick',0:0.2:1)
        if(i1==1)
            ylabel('disease prevalence','interpreter','latex','fontsize',16)
            title(strcat('$\beta=',num2str(beta),'$'),'interpreter','latex','fontsize',12)
        else
            title(strcat('$\alpha=',num2str(alpha),'$'),'interpreter','latex','fontsize',12)
        end
        text(-0.1,1.1,labs{i1},'fontsize',12)
        
        subplot(2,2,2+i1)
        hold on
        plot(r,diff1,'k')
        plot(diffpeaks(1),maxdiff1,'k*','MarkerSize', 5)
        plot(diffpeaks(2),abs(maxdiff2),'k*','MarkerSize', 5)
        ylim([0,0.5])
        if(i1==1)
            text(pos1(j1),maxdiff1+0.03,strcat('$\alpha=',num2str(alphaF),'$'),'interpreter','latex','fontsize',10)
        else
            text(pos2(j1),maxdiff1+0.03,strcat('$\beta=',num2str(betaF),'$'),'interpreter','latex','fontsize',10)
        end
        set(gca,'fontsize',8)
        box on
        set(gca,'ytick',0:0.1:0.5)
        if(i1==1)
            ylabel('$|D_M-D_F|$','interpreter','latex','fontsize',16)
            x1=xlabel('sex ratio at birth (proportion male), $r$','interpreter','latex','fontsize',16);
            temp = get(x1,'position');
            temp(1)=temp(1)+0.23;
            set(x1,'position',temp)
            title(strcat('$\beta=',num2str(beta),'$'),'interpreter','latex','fontsize',12)
        else
            title(strcat('$\alpha=',num2str(alpha),'$'),'interpreter','latex','fontsize',12)
        end
        text(-0.1,0.5*1.1,labs{i1+2},'fontsize',12)
    end
end
% save2pdf('fig1.pdf')
```
