## Supplementary material for "How do biases in sex ratio and disease characteristics affect the spread of sexually transmitted infections?": Source code for the model and figures: fig2.m

```
% Fig 2

alphaF = 0.3;
betaF = 0.5;

ALPHAM = [0.6,alphaF,0];
BETAM = [0.2,betaF,1];

r = linspace(0,1,101);
plotflag = 0;
c = 5;
d = 0.1;
sub = 0;

labs = {'(a)','(b)','(c)','(d)','(e)','(f)','(g)','(h)','(i)'};

str1 = {'>','=','<'};
str2 = {'<','=','>'};

figure(2)
clf
set(gcf,'color','w')
for i1=1:length(ALPHAM)
    alphaM = ALPHAM(i1);
    for j1=1:length(BETAM)
        betaM = BETAM(j1);
        
        males=zeros(length(r),1);
        females=zeros(length(r),1);
        overall=zeros(length(r),1);
        for i=1:length(r)
            [t,x] = sexratio1(alphaM,alphaF,betaM,betaF,r(i),plotflag);
            if(sum(x(end,3:4),2)>1e-6)
                males(i,1) = x(end,3)/sum(x(end,[1,3]),2);
                females(i,1) = x(end,4)/sum(x(end,[2,4]),2);
                overall(i,1) = sum(x(end,3:4),2)/sum(x(end,:),2);
            end
        end
        maxMP = max(males);
        maxFP = max(females);
        peaks = [r(find(males==max(males),1)),r(find(females==max(females),1))];
        
        sub=sub+1;
        subplot(length(ALPHAM),length(BETAM),sub)
        hold on
        plot(r,males,'k')
        plot(r,females,'k--')
        plot(r,overall,'k:','linewidth',1)
        plot(peaks(1),maxMP,'ko','MarkerSize', 5,'markerfacecolor','k')
        plot(peaks(2),maxFP,'ko','MarkerSize', 5,'markerfacecolor','w')
        ylim([0 1])
        box on
        set(gca,'fontsize',8)
        if(i1==2 && j1==1)
            ylabel('disease prevalence','interpreter','latex','fontsize',16)
        end
        if(i1==3 && j1==2)
            xlabel('sex ratio at birth (proportion male), $r$','interpreter','latex','fontsize',16)
        end
        text(-0.1,1.1,labs{sub},'fontsize',12)
        title(strcat('$\alpha_M',str1{i1},'\alpha_F$, $\beta_M',str2{j1},'\beta_F$'),'interpreter','latex','fontsize',12)
    end
end

save2pdf('fig2.pdf')
```
