## Supplementary material for "How do biases in sex ratio and disease characteristics affect the spread of sexually transmitted infections?": Source code for the model and figures: fig3.m

```
% Fig 3

clear
close all

format short

t_max = 2000;
dM = 0.1;
dF = dM;
gammaM = 0.2;
gammaF = 0.2;
b = 1;
f_F = 1;
f_M = 1;
q = 1e-3;
alphaM = 0.3;
alphaF = 0.3;
betaM = 0.5;
betaF = 0.5;
r = linspace(0,1,501);
betaMArray = linspace(0,1,500);
alphaMArray = linspace(0,2,500);
plotflag = 0;
c = 5;
d = 0.1;

malesbeta = zeros(length(betaMArray),length(r));
femalesbeta = zeros(length(betaMArray),length(r));
overallbeta = zeros(length(betaMArray),length(r));

% FIGURE 3(a-c)-------------------------------------------------------------
disp('Generating figure 3a-c')

for i=1:length(r)
    parfor j = 1 : length(betaMArray)
        [~,x] = sexratio1_fast(t_max,alphaM,alphaF,betaMArray(j),betaF,d,d,gamma,gamma,b,c,f_M,f_F,q,r(i));
        if(sum(x(end,3:4),2)>1e-6)
            malesbeta(j,i) = x(end,3)/sum(x(end,[1,3]),2);
            femalesbeta(j,i) = x(end,4)/sum(x(end,[2,4]),2);
            overallbeta(j,i) = sum(x(end,3:4),2)/sum(x(end,:),2);
        end
    end
    disp(strcat(num2str(i*100/length(r)),'% complete'));
end

malesalpha = zeros(length(alphaMArray),length(r));
femalesalpha = zeros(length(alphaMArray),length(r));
overallalpha = zeros(length(alphaMArray),length(r));


% FIGURE 3(d-f)-------------------------------------------------------------
disp('Generating figure 3d-f')

for i=1:length(r)
    parfor j = 1 : length(alphaMArray)
        [~,x] = sexratio1_fast(t_max,alphaMArray(j),alphaF,betaM,betaF,d,d,gamma,gamma,b,c,f_M,f_F,q,r(i));
        if(sum(x(end,3:4),2)>1e-6)
            malesalpha(j,i) = x(end,3)/sum(x(end,[1,3]),2);
            femalesalpha(j,i) = x(end,4)/sum(x(end,[2,4]),2);
            overallalpha(j,i) = sum(x(end,3:4),2)/sum(x(end,:),2);
        end
    end
    disp(strcat(num2str(i*100/length(r)),'% complete'));
end

save('fig3.mat')
%%
load('fig3.mat')
malesalpha(malesalpha==0)=NaN;
femalesalpha(femalesalpha==0)=NaN;
overallalpha(overallalpha==0)=NaN;
malesbeta(malesbeta==0)=NaN;
femalesbeta(femalesbeta==0)=NaN;
overallbeta(overallbeta==0)=NaN;

figure(3)
clf
set(gcf,'color','w')
labs = {'(a)','(b)','(c)','(d)','(e)','(f)'};

subplot(2,3,1)
contourf(r,betaMArray,malesbeta);
hold on
plot([0.5,0.5],[0,1],'r','linewidth',2)
plot([0,1],[betaF,betaF],'r--','linewidth',2)
set(gca,'clim',[0,0.7])
title('Males','interpreter','latex','fontsize',12);
text(0.01,0.05,'$R_0<1$','fontsize',10,'interpreter','latex');    
ylabel({'Female-to-male','transmission rate, $\beta_{M}$'},'interpreter','latex','fontsize',14);

subplot(2,3,2)
contourf(r,betaMArray,femalesbeta);
hold on
plot([0.5,0.5],[0,1],'r','linewidth',2)
plot([0,1],[betaF,betaF],'r--','linewidth',2)
set(gca,'clim',[0,0.7])
title('Females','interpreter','latex','fontsize',12);
text(0.01,0.05,'$R_0<1$','fontsize',10,'interpreter','latex');    

subplot(2,3,3)
contourf(r,betaMArray,overallbeta);
hold on
plot([0.5,0.5],[0,1],'r','linewidth',2)
plot([0,1],[betaF,betaF],'r--','linewidth',2)
set(gca,'clim',[0,0.7])
temp1 = get(gca,'position');
title('Overall','interpreter','latex','fontsize',12);
text(0.01,0.05,'$R_0<1$','fontsize',10,'interpreter','latex');    

subplot(2,3,4)
contourf(r,alphaMArray,malesalpha);
hold on
plot([0.5,0.5],[0,2],'r','linewidth',2)
plot([0,1],[alphaF,alphaF],'r--','linewidth',2)
set(gca,'clim',[0,0.7])
title('Males','interpreter','latex','fontsize',12);
text(0.01,1.88,'$R_0<1$','fontsize',10,'interpreter','latex');    
xlim([0,1])
ylabel({'Mortality virulence','in males, $\alpha_{M}$'},'interpreter','latex','fontsize',14);

subplot(2,3,5)
contourf(r,alphaMArray,femalesalpha);
hold on
plot([0.5,0.5],[0,2],'r','linewidth',2)
plot([0,1],[alphaF,alphaF],'r--','linewidth',2)
xlabel('Sex ratio at birth (proportion male), $r$','interpreter','latex','fontsize',14);
set(gca,'clim',[0,0.7])
title('Females','interpreter','latex','fontsize',12);
text(0.01,1.88,'$R_0<1$','fontsize',10,'interpreter','latex');    
xlim([0,1])

subplot(2,3,6)
contourf(r,alphaMArray,overallalpha);
hold on
plot([0.5,0.5],[0,2],'r','linewidth',2)
plot([0,1],[alphaF,alphaF],'r--','linewidth',2)
set(gca,'clim',[0,0.7])
temp2 = get(gca,'position');
title('Overall','interpreter','latex','fontsize',12);
text(0.01,1.88,'$R_0<1$','fontsize',10,'interpreter','latex');    
xlim([0,1])
drawnow

for i=1:3
    subplot(2,3,i)
    temp = get(gca,'position');
    temp(1) = temp(1)-0.02;
    set(gca,'position',temp);
    text(0,1.1,labs{i},'fontsize',12);
end
for i=4:6
    subplot(2,3,i)
    temp = get(gca,'position');
    temp(1) = temp(1)-0.02;
    set(gca,'position',temp);
    text(0,2.2,labs{i},'fontsize',12);
end

C = colorbar;
Cpos = [temp1(1)+temp1(3),temp2(2),0.02,temp1(4)+temp1(2)-temp2(2)-0.013];
set(C,'position',Cpos);
y1=ylabel(C,'disease prevalence','interpreter','latex','fontsize',14);
load('cvidis.mat')
colormap(cvidis);

% save2pdf('fig3.pdf')
```
