## Supplementary material for "How do biases in sex ratio and disease characteristics affect the spread of sexually transmitted infections?": Source code for the model and figures: fig4.m

```
% Fig 4

clear
close all

format short

t_max = 2000;
alphaM = 0.3;
alphaF = 0.3;
betaM = 0.5;
betaF = 0.5;
r = linspace(0,1,101);
betaMArray = linspace(0,1,501);
alphaMArray = linspace(0,2,500);
d = 0.1;
gammaM = 0.2;
gammaF = 0.2;
b = 1;
c = 5;
f_F = 1;
f_M = 1;
q = 1e-3;
d = 0.1;
plotflag = 0;

malespeak = zeros(length(betaMArray),length(alphaMArray));
femalespeak = zeros(length(betaMArray),length(alphaMArray));
overallpeak = zeros(length(betaMArray),length(alphaMArray));

% FIGURE 4 -------------------------------------------------------------

disp('Generating figure 4');

for i=1:length(betaMArray)
    for j = 1:length(alphaMArray)
        males = zeros(1,length(r));
        females = zeros(1,length(r));
        overall = zeros(1,length(r));
        parfor k = 1:length(r)
            [~,x] = sexratio1_fast(t_max,alphaMArray(j),alphaF,betaMArray(i),betaF,d,d,gammaM,gammaF,b,c,f_M,f_F,q,r(k));
            if(sum(x(end,3:4),2)>1e-6)
                males(1,k) = x(end,3)/sum(x(end,[1,3]),2);
                females(1,k) = x(end,4)/sum(x(end,[2,4]),2);
                overall(1,k) = sum(x(end,3:4),2)/sum(x(end,:),2);
            end
        end
        males = smooth(males,10);
        females = smooth(females,10);
        overall = smooth(overall,10);
        malespeak(i,j) = r(find(males==max(males),1));
        femalespeak(i,j) = r(find(females==max(females),1));
        overallpeak(i,j) = r(find(overall==max(overall),1));
    end
    disp(strcat(num2str(i*100/length(betaMArray)),'% complete'));
end
clear i j k t x ans males females overall

save('fig4.mat')
%%
load('fig4.mat')

malespeak(malespeak==0)=NaN;
femalespeak(femalespeak==0)=NaN;
overallpeak(overallpeak==0)=NaN;

mn = min([min(malespeak),min(femalespeak),min(overallpeak(:))]);
mx = max([max(malespeak),max(femalespeak),max(overallpeak(:))]);

mn = floor(mn*10)/10;
mx = ceil(mx*10)/10;

figure(4)
clf
set(gcf,'color','w')

subplot(1,3,1)
hold on
contourf(alphaMArray,betaMArray,malespeak);set(gca,'clim',[mn,mx])
plot([0,2],[0.5,0.5],'r--','linewidth',2)
plot([alphaF,alphaF],[0,1],'r','linewidth',2)
box on
title('Males','interpreter','latex','fontsize',12);
ylabel({'Female-to-male','transmission rate, $\beta_{M}$'},'interpreter','latex','fontsize',16);
text(0,1.09,'(a)','fontsize',12);

subplot(1,3,2)
hold on
contourf(alphaMArray,betaMArray,femalespeak);set(gca,'clim',[mn,mx])
plot([0,2],[0.5,0.5],'r--','linewidth',2)
plot([alphaF,alphaF],[0,1],'r','linewidth',2)
box on
title('Females','interpreter','latex','fontsize',12);
xlabel('Mortality virulence in males, $\alpha_{M}$','interpreter','latex','fontsize',16);
text(0,1.09,'(b)','fontsize',12);

subplot(1,3,3)
hold on
contourf(alphaMArray,betaMArray,overallpeak);set(gca,'clim',[mn,mx])
plot([0,2],[0.5,0.5],'r--','linewidth',2)
plot([alphaF,alphaF],[0,1],'r','linewidth',2)
box on
title('Overall','interpreter','latex','fontsize',12);
text(0,1.09,'(c)','fontsize',12);

for i=1:3
    subplot(1,3,i)
    temp = get(gca,'position');
    temp(1)=temp(1)-0.03;
    temp(2)=temp(2)+0.08;
    temp(4)=temp(4)-0.1;
    set(gca,'position',temp)
    text(1.2,0.12,'$R_0<1$','fontsize',12,'interpreter','latex');
end
drawnow

C = colorbar;
Cpos = get(C,'position');
Cpos(1) = temp(1)+temp(3)+0.02;
set(C,'position',Cpos);
y1=ylabel(C,{'Sex ratio at birth where','disease prevalence peaks'},'interpreter','latex','fontsize',13);
set(C,'ytick',mn:0.1:mx)

load('cvidis.mat')
colormap(cvidis);

% fig4=figure(4);
% fig4.Renderer='Painters';
% save2pdf('fig4.pdf')
```
