## Supplementary material for "How do biases in sex ratio and disease characteristics affect the spread of sexually transmitted infections?": Source code for the model and figures: sexratio1.m

```
function [t,x] = sexratio1(alphaM,alphaF,betaM,betaF,r,plotflag)

t_max = 2000;
%betaM = beta;
%betaF = beta;
deltaM = 0.1;
deltaF = deltaM;
gammaM = 0.2;
gammaF = 0.2;
c = 5;
b = 1;
f_F = 1;
f_M = 1;
q = 1e-3;

init_pop = [r,(1-r),r/10,(1-r)/10]; % Initial population [SM,SF,IM,IF]

pars = [];

% All checks completed, now solve the equations
[t,x] = ode45(@ode,[0,t_max],init_pop,odeset('abstol',1e-10,'nonnegative',1:length(init_pop)),pars);

    function dxdt = ode(t,x,~)
        
        SM = x(1);
        SF = x(2);
        IM = x(3);
        IF = x(4);
        N = sum(x);
        
        SMSF = c*SM*SF/N;
        SMIF = c*SM*IF/N;
        IMSF = c*IM*SF/N;
        IMIF = c*IM*IF/N;
        
        births = b*(1-q*N)*(SMSF + f_F*SMIF + f_M*IMSF + f_M*f_F*IMIF);
        
        dxdt = [births*r - deltaM*SM - betaM*SMIF + gammaM*IM;
            births*(1-r) - deltaF*SF - betaF*IMSF + gammaF*IF;
            betaM*SMIF - (alphaM + deltaM + gammaM)*IM;
            betaF*IMSF - (alphaF + deltaF + gammaF)*IF];
    end

if(plotflag>0)
    figure(2)
    clf
    hold on
    plot(t,x(:,3)./sum(x(:,[1,3]),2),'b')
    plot(t,x(:,4)./sum(x(:,[2,4]),2),'r')
    plot(t,sum(x(:,3:4),2)./sum(x,2),'k')
    hold off
    legend('male','female','overall')
    xlabel('Time')
    ylabel('disease prevalence')
    
    figure(3)
    clf
    hold on
    plot(t,x(:,3),'b')
    plot(t,x(:,4),'r')
    plot(t,sum(x(:,3:4),2),'k')
    hold off
    legend('male','female','overall')
    xlabel('Time')
    ylabel('Infected individuals')
end

end
```
