## Supplementary material for "How do biases in sex ratio and disease characteristics affect the spread of sexually transmitted infections?": Source code for the model and figures: sexratio1_fast.c

```
/*****************************************************************************************************
 * [t,x] = sexratio1_fast(t_max,alphaM,alphaF,betaM,betaF,dM,dF,gammaM,gammaF,b,c,f_M,f_F,q,r)
 *
 * Compile in Matlab using mex sexratio1_fast.c
 ****************************************************************************************************/

#include <mex.h>
#include <math.h>

/***********************************
 * CONSTANT PARAMATER VALUES
 ***********************************/
#define MAXSTEPS 1e6 /* Maximum number of steps for ODP solver */
#define EPS 1e-3 /* ODE solver tolerance */
#define TINY 1e-30 /* Constant value for solver */
#define b21 0.2
#define b31 3.0/40.0
#define b32 9.0/40.0
#define b41 0.3
#define b42 -0.9
#define b43 1.2
#define b51 -11.0/54.0
#define b52 2.5
#define b53 -70.0/27.0
#define b54 35.0/27.0
#define b61 1631.0/55296
#define b62 175.0/512.0
#define b63 575.0/13824.0
#define b64 44275.0/110592
#define b65 253.0/4096.0
#define c1 37.0/378.0
#define c3 250.0/621.0
#define c4 125.0/594.0
#define c6 512.0/1771.0
#define dc5 -277.00/14336
// #define isnan(x) ((x) != (x))

struct PARAM{
    double t_max;
    double alphaM;
    double alphaF;
    double betaM;
    double betaF;
    double dM;
    double dF;
    double gammaM;
    double gammaF;
    double b;
    double c;
    double f_M;
    double f_F;
    double q;
    double r; 
};

/*************************************
 * Function prototypes
 *************************************/
int my_rungkut (double *T, double *xOut, struct PARAM *p);
void rkqs(double *x, double *dxdt, double *h, double *hnext, double *xScale, struct PARAM *p);
void rkck(double *x, double *dxdt, double *xOut, double *xErr, double h, struct PARAM *p);
void dynamic(double *x, double *dxdt, struct PARAM *p);
double FMAX(double, double);
double FMIN(double, double);

void mexFunction(int nlhs, mxArray *plhs[], int nrhs, const mxArray *prhs[])
{
    double *t, *x, *parameter, *tTemp, *xTemp;
    int i, j, colLen, maxsteps;
    struct PARAM p;
    
    /* Allocate inputs */
    if(nrhs!=15){
        mexErrMsgTxt("Incorrect number of input arguments!\n");
    }
    else{
        parameter= mxGetPr(prhs[0]);
        p.t_max= *parameter;
        parameter= mxGetPr(prhs[1]);
        p.alphaM= *parameter;
        parameter= mxGetPr(prhs[2]);
        p.alphaF= *parameter;
        parameter= mxGetPr(prhs[3]);
        p.betaM= *parameter;
        parameter= mxGetPr(prhs[4]);
        p.betaF= *parameter;
        parameter= mxGetPr(prhs[5]);
        p.dM= *parameter;
        parameter= mxGetPr(prhs[6]);
        p.dF= *parameter;
        parameter= mxGetPr(prhs[7]);
        p.gammaM= *parameter;
        parameter= mxGetPr(prhs[8]);
        p.gammaF= *parameter;
        parameter= mxGetPr(prhs[9]);
        p.b= *parameter;
        parameter= mxGetPr(prhs[10]);
        p.c= *parameter;
        parameter= mxGetPr(prhs[11]);
        p.f_M= *parameter;
        parameter= mxGetPr(prhs[12]);
        p.f_F= *parameter;
        parameter= mxGetPr(prhs[13]);
        p.q= *parameter;
        parameter= mxGetPr(prhs[14]);
        p.r= *parameter;
    }
    maxsteps = (int)MAXSTEPS;
    
    /* Allocate memory */
    tTemp = malloc(maxsteps*sizeof(double));
    xTemp = malloc(maxsteps*4*sizeof(double));
    
    /* Call ODP solver */
    colLen = my_rungkut(tTemp, xTemp, &p);
    
    /* Create outputs */
    plhs[0] = mxCreateDoubleMatrix(colLen, 1, mxREAL);
    plhs[1] = mxCreateDoubleMatrix(colLen, 4, mxREAL);
    
    t = mxGetPr(plhs[0]);
    x = mxGetPr(plhs[1]);
    
    /* Copy data to outputs */
    for (i=0;i<colLen;i++){
        t[i] = tTemp[i];
        for (j=0;j<4;j++) {
            x[i + j*colLen] = xTemp[i + j*maxsteps];
        }
    }
    
    /* Free memory */
    free(tTemp);
    free(xTemp);
    
    return;
}

/*****************************************
 * ODP solver
 ****************************************/
int my_rungkut (double *T, double *xOut, struct PARAM *p){
    
    double t, x[4], dxdt[4], xScale[4], hnext[1], h[1];
    int i, j, k, exitflag, count, maxsteps;
    
    /* Other parameters */
    exitflag = 1;
    count=0;
    k=1;
    h[0] = 1e-3;
    hnext[0] = 1e-3;
    t=0;
    maxsteps = (int)MAXSTEPS;
    
    /* Initialise populations */
    x[0] = p->r;
    x[1] = 1-p->r;
    x[2] = p->r/10;
    x[3] = (1-p->r)/10;
    
    /* Update output */
    T[0]=t;
    for (i=0; i<4; i++) {
        xOut[i*maxsteps] = x[i];
    }
    
    /* Main loop: */
    do{
        /* This ensures the final step lands us on the final time point */
        if(1.1*hnext[0]>(p->t_max-t)){
            hnext[0] = p->t_max-t;
            h[0] = p->t_max-t;
            t=p->t_max;
            exitflag=0;
        }
        else{
            h[0] = hnext[0];
            t+=h[0];
        }
        if(t>=p->t_max) {
            t=p->t_max;
            exitflag=0;
        }
        /* This is where the equations are first solved */
        dynamic(x, dxdt, p);

        /* Adjust the step size to maintain accuracy */
        for (i=0; i<4; i++){
            x[i] = FMAX(x[i],0);
            xScale[i]=fabs(x[i])+fabs(dxdt[i]*(*h))+TINY;
        }
        
        rkqs(x, dxdt, h, hnext, xScale, p);
        
        for (i=0; i<4; i++){
            x[i] = FMAX(x[i],0);
        }
        
        /* Update output */
        count++;
        T[count] = t;
        for (i=0; i<4; i++) {
            xOut[count + i*maxsteps] = x[i];
        }
    }while(count<(maxsteps-1) && t<=p->t_max && exitflag);
    count++;
    
    return count;
}

/***************************************
 * This generates the adaptive step-size
 **************************************/
void rkqs(double *x, double *dxdt, double *h, double *hnext, double *xScale, struct PARAM *p)
{
    double xTemp[4], xErr[4], htemp, errmax;
    int i, j, count;
    
    count = 0;
    while(count<1e5)
    {
        
        rkck(x, dxdt, xTemp, xErr, *h, p);
        
        errmax= 0.0;
        for(i=0;i<4;i++){
            errmax= FMAX(errmax, fabs(xErr[i]/(xScale[i])));
        }
        errmax/= EPS;
        if(errmax<=1.0) break;
        htemp= 0.9*(*h)*pow(errmax, -0.25);
        *h= (*h>=0.0 ? FMAX(htemp, 0.1*(*h)) : FMIN(htemp, 0.1*(*h)));
        count++;
        if(count>1e4){
            mexErrMsgTxt("stuck in loop!\n");
            break;
        }
    }    
    if(errmax > 1.89E-4) {
        *hnext= 0.9*(*h)*pow(errmax, -0.2);
    }
    else {
        *hnext= 5.0*(*h);
    }
    
    for(i=0;i<4;i++){
        x[i] = xTemp[i];
    }
}

/**************************************
 * Standard RK solver
 **************************************/
void rkck(double *x, double *dxdt, double *xOut, double *xErr, double h, struct PARAM *p){
    
    int i, j;
    double xk1[4], xk2[4], xk3[4], xk4[4], xk5[4], xk6[4], xTemp[4];
    double dc1=c1-2825.0/27648.0, dc3=c3-18575.0/48384.0, dc4=c4-13525.0/55296.0,
            dc6=c6-0.25;
    
    for(i=0;i<4;i++){
        xTemp[i] = x[i] + b21*h*dxdt[i];
    }
    dynamic(xTemp, xk2, p);
    
    for(i=0;i<4;i++){
        xTemp[i] = x[i]+h*(b31*dxdt[i]+b32*xk2[i]);
    }
    dynamic(xTemp, xk3, p);
    
    for(i=0;i<4;i++){
        xTemp[i] = x[i]+h*(b41*dxdt[i]+b42*xk2[i]+b43*xk3[i]);
    }
    dynamic(xTemp, xk4, p);
    
    for(i=0;i<4;i++){
        xTemp[i] = x[i]+h*(b51*dxdt[i]+b52*xk2[i]+b53*xk3[i]+b54*xk4[i]);
    }
    dynamic(xTemp, xk5, p);
    
    for(i=0;i<4;i++){
        xTemp[i] = x[i]+h*(b61*dxdt[i]+b62*xk2[i]+b63*xk3[i]+b64*xk4[i]+b65*xk5[i]);
    }
    dynamic(xTemp, xk6, p);
    
    for(i=0;i<4;i++){
        xOut[i]= x[i]+h*(c1*dxdt[i]+c3*xk3[i]+c4*xk4[i]+c6*xk6[i]);
        xErr[i]= h*(dc1*dxdt[i]+dc3*xk3[i]+dc4*xk4[i]+dc5*xk5[i]+dc6*xk6[i]);
    }
}

/**************************************
 * Population and evolutionary dynamics
 **************************************/
void dynamic(double *x, double *dxdt, struct PARAM *p){
    
    int i, j;
    double SM, SF, IM, IF, N, SMSF, SMIF, IMSF, IMIF, births;
   
    SM = x[0];
    SF = x[1];
    IM = x[2];
    IF = x[3];
    N = SM + SF + IM + IF;
        
    SMSF = p->c*SM*SF/FMAX(TINY,N);
    SMIF = p->c*SM*IF/FMAX(TINY,N);
    IMSF = p->c*IM*SF/FMAX(TINY,N);
    IMIF = p->c*IM*IF/FMAX(TINY,N);
        
    births = p->b*(1-p->q*N)*(SMSF + p->f_F*SMIF + p->f_M*IMSF + p->f_M*p->f_F*IMIF);
    
    dxdt[0] = births*p->r - p->dM*SM - p->betaM*SMIF + p->gammaM*IM;
    dxdt[1] = births*(1-p->r) - p->dF*SF - p->betaF*IMSF + p->gammaF*IF;
    dxdt[2] = p->betaM*SMIF - (p->alphaM + p->dM + p->gammaM)*IM;
    dxdt[3] = p->betaF*IMSF - (p->alphaF + p->dF + p->gammaF)*IF;
}

/***************************************
 * Return maximum of two inputs
 ***************************************/
double FMAX(double l, double r)
{
    if(l>r)return l;
    else   return r;
}

/***************************************
 * Return minimum of two inputs
 ***************************************/
double FMIN(double l, double r)
{
    if(l<r)return l;
    else   return r;
}
```
